# Spatiospectral image processing workflow considerations for advanced MR spectroscopy of the brain

**DOI:** 10.1101/2023.09.07.556701

**Authors:** Leon Y. Cai, Stephanie N. Del Tufo, Laura Barquero, Micah D’Archangel, Lanier Sachs, Laurie E. Cutting, Nicole Glaser, Simona Ghetti, Sarah S. Jaser, Adam W. Anderson, Lori C. Jordan, Bennett A. Landman

## Abstract

Magnetic resonance spectroscopy (MRS) is one of the few non-invasive imaging modalities capable of making neurochemical and metabolic measurements *in vivo*. Traditionally, the clinical utility of MRS has been narrow. The most common use has been the “single-voxel spectroscopy” variant to discern the presence of a lactate peak in the spectra in one location in the brain, typically to evaluate for ischemia in neonates. Thus, the reduction of rich spectral data to a binary variable has not classically necessitated much signal processing. However, scanners have become more powerful and MRS sequences more advanced, increasing data complexity and adding 2 to 3 spatial dimensions in addition to the spectral one. The result is a spatially- and spectrally-variant MRS image ripe for image processing innovation. Despite this potential, the logistics for robustly accessing and manipulating MRS data across different scanners, data formats, and software standards remain unclear. Thus, as research into MRS advances, there is a clear need to better characterize its image processing considerations to facilitate innovation from scientists and engineers. Building on established neuroimaging standards, we describe a framework for manipulating these images that generalizes to the voxel, spectral, and metabolite level across space and multiple imaging sites while integrating with LCModel, a widely used quantitative MRS peak-fitting platform. In doing so, we provide examples to demonstrate the advantages of such a workflow in relation to recent publications and with new data. Overall, we hope our characterizations will lower the barrier of entry to MRS processing for neuroimaging researchers.

## 1. INTRODUCTION

Magnetic resonance imaging (MRI) has long been an established imaging modality for assessing the *structure* of the brain. However, one key limitation is that it does not provide any information about *neurochemical* phenomena in the brain. MR spectroscopy (MRS), however, fills this gap. By changing how information is encoded in MRI, MRS sequences allow scanners to capture local magnetic field changes due to the chemical environments surrounding protons, thus assessing neurochemical phenomena and, by extension, metabolic ones [1], [2]. This information is encoded in chemical shifts of the resonant frequency of protons in the spectral, or frequency, domain [1], [2]. Thus, MRS data in a voxel consists not of a single intensity, as in typical MRI, but of a spectrum of peaks, each one corresponding to protons from different local chemical environments. By assessing the peaks, one can therefore gain insight about the underlying metabolites.

MRS generally is one of two types: single-voxel spectroscopy (SVS) and chemical shift imaging (CSI). In the case of SVS, data are acquired in one voxel (i.e., no spatial dimension) as opposed to in a 2-dimensional spatial slice of voxels or 3-dimensional spatial volume of voxels for CSI. These spectral peaks are then analyzed to discern the presence of certain metabolites [1], [2]. In the brain, these include lactate, N-acetylaspartate (NAA), and creatine (Cr), among many others.

The way these spectra are analyzed, however, varies by application. For instance, clinically, one of the most common uses of MRS in the brain is as SVS in neonates to evaluate for ischemic encephalopathy [3]. In these cases, a spectrum is acquired in a single voxel, typically in a deep gray matter nucleus, like the basal ganglia. The spectrum is then subsequently visualized, and the presence of an elevated lactate peak is discerned *qualitatively* (Figure 1a) [4]. As lactate is considered to be a marker for ischemia, this information provides an additional data point for clinicians, but effectively ignores the rich information available in the remainder of the spectra and fails to provide *quantitative* estimates of the size of the lactate peak [4]. Further, as SVS contains no spatial dimensions, this visual approach effectively reduces all information to a single binary variable: is lactate elevated or not in the basal ganglia?

**Figure 1.**
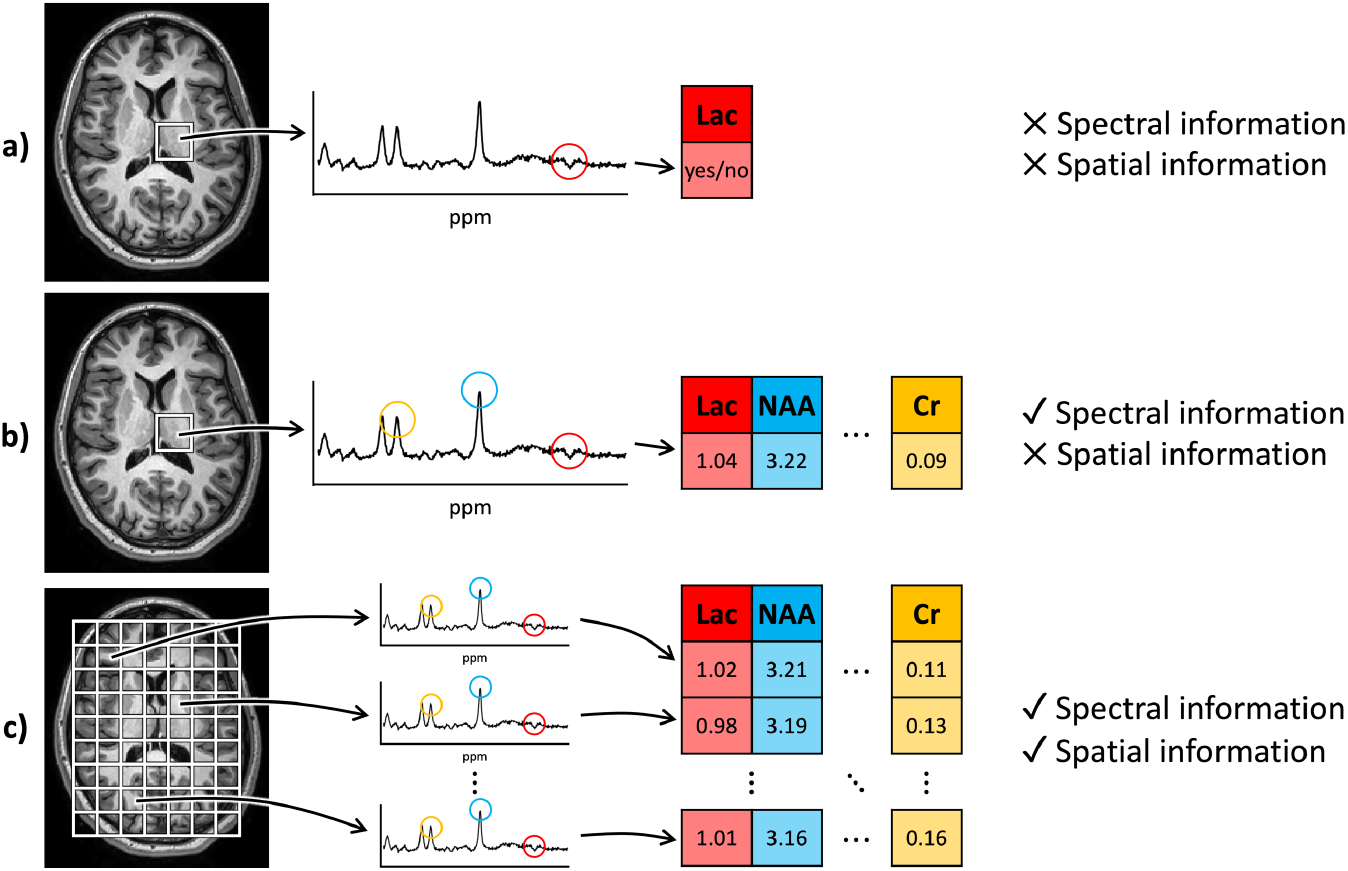
Overview of MRS approaches. (a) Clinically, SVS spectra are often visually reduced to a binary variable representing the presence or absence of a single metabolite peak. This approach does not take advantage of either spectral (or metabolite) or spatial information. (b) In research, SVS is used to quantify different metabolites in a single location, thus adding spectral information but lacking spatial information. (c) Up-and-coming CSI sequences are used in research to quantify spectral metabolite information at different locations, providing the opportunity for rich spatiospectral processing and inference. Lac = lactate.

In the research realm, however, SVS often does not suffer from the same limitations. As opposed to a qualitative analysis, a quantitative analysis of spectra, typically with peak fitting software like the industry standard LCModel, estimates the area under each of the peaks as a surrogate for the concentrations of the corresponding metabolites (Figure 1b) [5]. This approach avoids the reduction of spectral information to a single binary variable, but still does not provide any spatial data across different brain structures or regions. The addition of CSI, however, allows this approach to be repeated across the brain, thus providing a spatiospectrally varying image and enabling the simultaneous investigation of metabolites in different regions (Figure 1c) [6].

As such, MRS provides unique insight into the physiology of the brain, and innovations in CSI and spectral software contain rich potential for researchers to extract such information. That being said, the image processing landscape regarding MRS has yet to flourish. Unlike other MRI modalities like T1-weighted (T1w) MRI, functional MRI (fMRI), or diffusion MRI (dMRI) which have an established gamut of image processing tools and standards to improve data interpretation, MRS does not. This is largely due to the fragmented workflow and software landscape available to researchers, raising the barrier for existing neuroimaging investigators to make the jump from MRI to MRS. Thus, here we describe considerations taken by recent studies to improve the access of MRS to traditional neuroimaging researchers and consolidate them into a proposed workflow for future MRS studies [7].

## 2. METHODS

### 2.1 Existing and proposed workflows

In order to provide context for our proposed workflow, we first describe common workflows for other MRI modalities, notably fMRI and dMRI. Unlike traditional MRI which provides intensity information in 3-dimensional space, fMRI and dMRI contain an additional dimension: fMRI contains blood flow information across time and dMRI contains information regarding the movement of water across different directions. Thus, both fMRI and dMRI images are 4-dimensional.

To accommodate these data, a common fMRI and dMRI workflow has emerged that consists of three main steps (Figure 2a). The first is the exporting of raw data off the MRI scanner in DICOM format. Variability of this format between different scanner manufacturers and between different styles of DICOM files has subsequently led to the development of a common neuroimaging research standard, the NIFTI file. This file type has two main parts. The first is an array of numbers containing the image data, and the second is a header containing some of the imaging sequence parameters and more importantly an affine transform describing the spatial location of the array. With these two components, the NIFTI file format has spurred the development of numerous spatiotemporal (for fMRI) or spatial and gradient-based (for dMRI) image processing routines. In order to store fMRI-or dMRI-specific sequence parameters, the NIFTI paradigm typically uses affiliated text files or JSON sidecars for ease of interrogation. For instance, the gradient direction information for dMRI is stored in .bval and .bvec text files of length equivalent to the number of dMRI volumes whereas the phase-encoding information needed for distortion correction is often stored in a JSON sidecar [8], [9].

**Figure 2.**
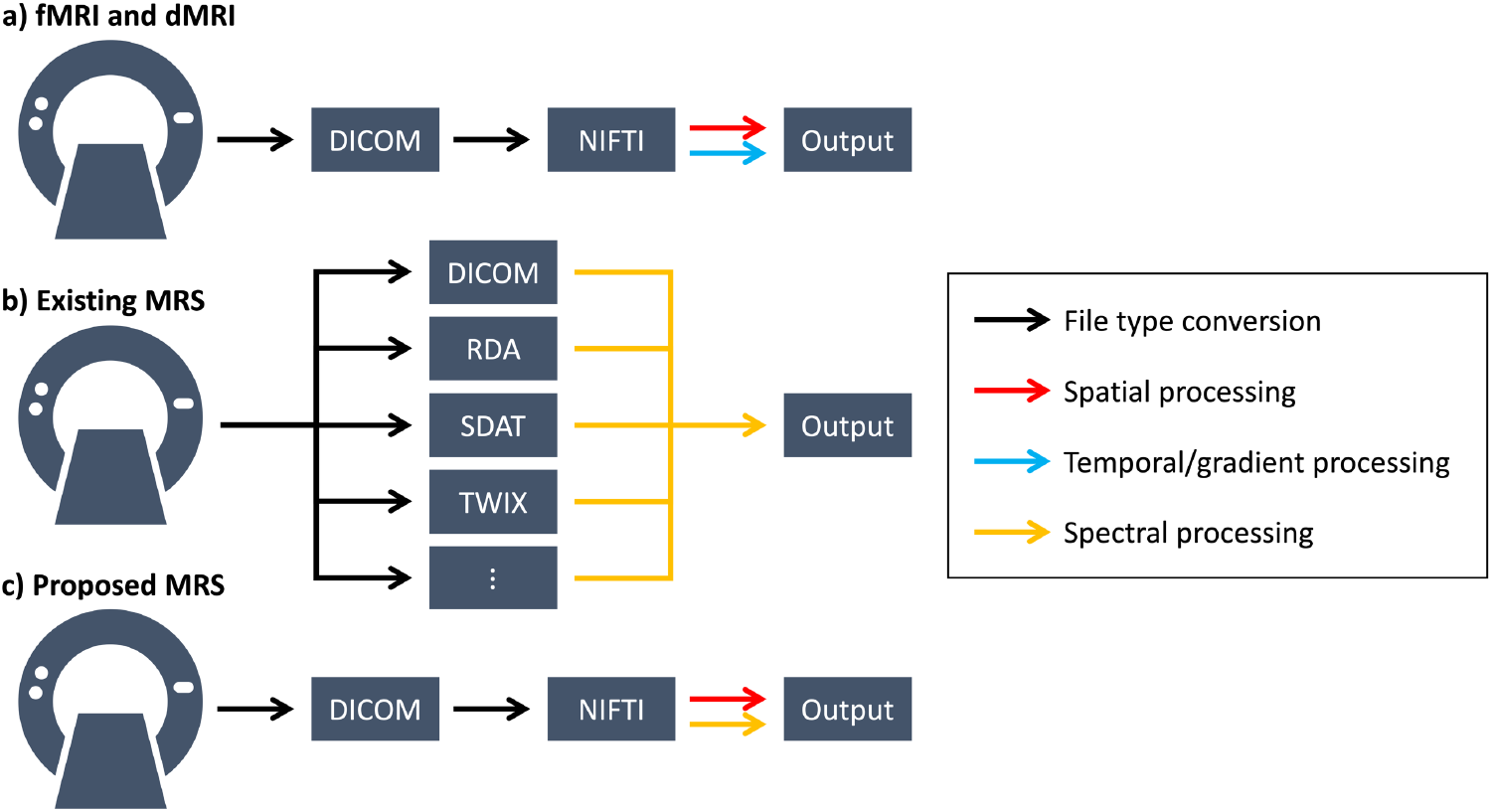
Existing and proposed workflows. (a) Existing fMRI and dMRI workflows rely on the NIFTI file standard to remove variability in DICOMs and facilitate spatiotemporal or spatial with gradient processing, respectively. (b) The fragmented data landscape for MRS has come to support spectral processing, but at the cost of increased logistical variability and minimal spatial processing support. (c) The proposed approach aligns MRS workflows with existing neuroimaging frameworks, replacing the temporal/gradient dimension with a spectral one, to facilitate spatiospectral processing and simplify entry into MRS analysis for those familiar with traditional MRI standards.

MRS, on the other hand, has not achieved such a standardized workflow (Figure 2b). For instance, in addition to the already variable DICOM format, some scanners can also export MRS data to proprietary formats like the RDA, SDAT, and TWIX file types. Fortunately, existing spectral processing software, like LCModel, supports many of these file types [5]. Unfortunately, however, many of these file types do not explicitly consider the spatial location and orientation of the spectral data array, thus making it difficult to implement spatial processing routines for CSI, analyze MRS images jointly with other MRI modalities already in the NIFTI format, and visualize MRS data without specialized software.

Thus, we propose consolidating the MRS workflow to match that of fMRI and dMRI, using NIFTI as a standard to save both the spectral data array and the spatial location and orientation transforms to facilitate spatiospectral processing and viewing of the data with existing NIFTI viewers (Figure 2c). Additionally, through associated text files or JSON sidecars, the unique sequence parameters for MRS can also be saved and interrogated [10]. These considerations allow MRS to be treated like any other neuroimaging modality and provide an intuitive framework for new users familiar with traditional neuroimaging workflows. This perspective aligns with the recent development of tools to facilitate the conversion of MRS DICOMs and other file types to NIFTI-based formats [11].

### 2.2 Data formatting considerations

One consideration for this proposed workflow is that like with other MRI modalities, the DICOM files produced by different scanner manufacturers can differ in their structure and data formatting. For instance, Philips MRS DICOMs contain two sets of data, a water-suppressed and water-unsuppressed spectrum (Figure 3a). These data reflect the need for MRS sequences to suppress the overwhelming water signal in order to capture chemical shifts in less abundant metabolites [2]. In Siemens MRS DICOMs, however, only one suppressed spectrum is made available from the scanner (Figure 3b). We note that researchers using Philips data must be careful in checking which of the two they are using when performing processing on the spectra themselves.

**Figure 3.**
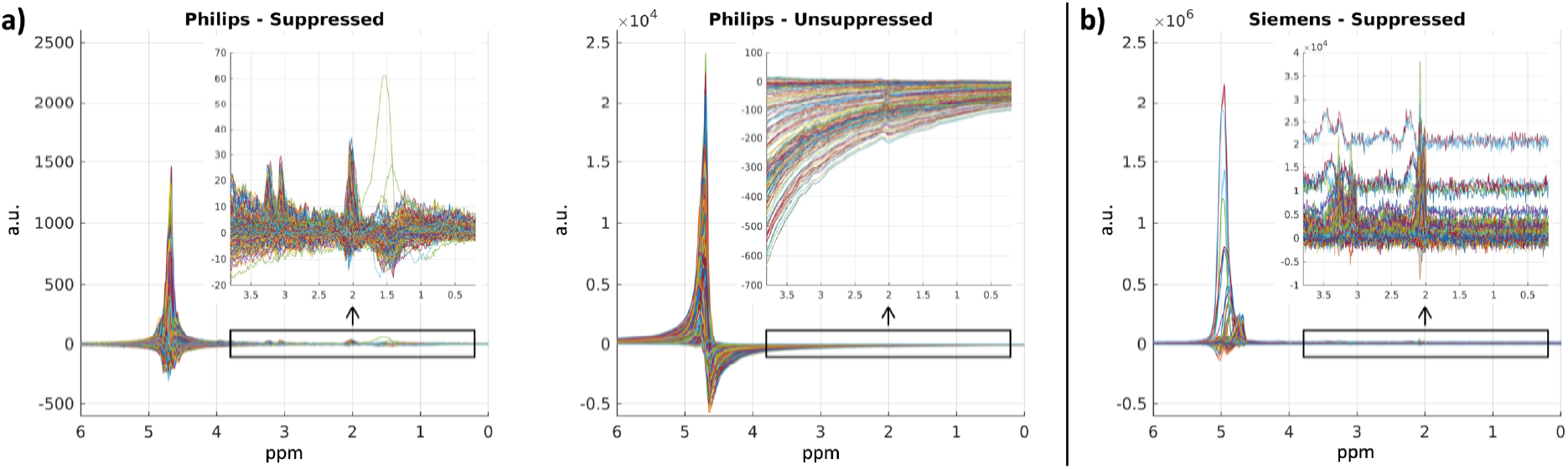
Variability in MRS data between scanners. As seen in the plotted real components of the complex-valued spectra, Philips MRS DICOMs contain both water-suppressed and water-unsuppressed data (a) whereas Siemens DICOMs contain only suppressed (b). Further, the signal amplitude between suppressed spectra can vary widely between scanner manufacturers. a.u. = arbitrary units.

Further, both Philips and Siemens use non-standard DICOM fields to encode and save their MRS data, which are provided as complex-valued time-domain free induction decay signals, as opposed to spectra in the Fourier domain. Fortunately, with standardization in the NIFTI format, converting between the two domains can be as simple as using the fast Fourier transform (FFT) and its inverse as implemented in existing software libraries like SciPy or MATLAB (MathWorks) along the spectral dimension of the array [12]. Of note, FFT typically outputs in Hz, whereas MRS is typically analyzed in ppm. We propose storing the necessary sequence parameters for Hz to ppm conversion in an associated text file and additionally save the ppm itself in a .ppm text file analogous to the .bval and .bvec dMRI files.

In addition to time-versus spectral-domain manipulation of the imaging array, another key consideration of conversion to NIFTI is the accuracy of the spatial orientation information in the header. Due to the low resolution of CSI (i.e., on the order of cm) compared to other MRI modalities (i.e., on the order of mm), discerning whether an image is flipped, rotated, or transposed can be difficult (Figure 4a). Thus, we recommend the use of coarse phantom studies to ensure images are oriented properly after conversion to NIFTI (Figure 4b). For Figure 4, we achieve this using water bottles and the Braino MRS phantom (GE Healthcare) placed in the scanner.

**Figure 4.**
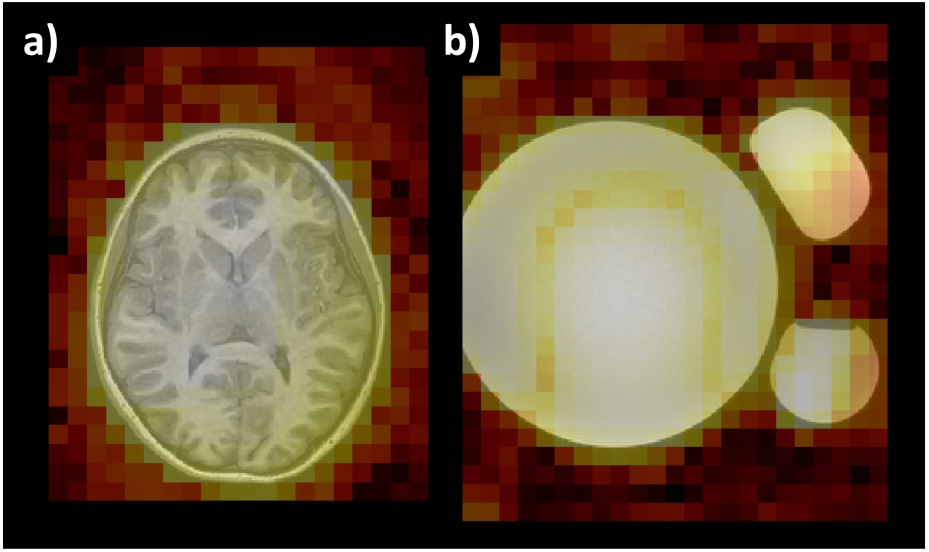
The low resolution of CSI necessitates verification of spatial orientation. (a) Brain anatomy is largely left-right symmetric, especially when the resolution is on the order of cm with CSI, making it difficult to discern orientation accuracy. (b) Coarse asymmetric phantom studies can help ensure DICOM to NIFTI conversions are accurate. The MRS heat maps are produced by collapsing the spectral dimension across ppm with the log average real amplitude.

### 2.3 Data collection

To demonstrate the utility of the proposed workflow across various types of studies, we used data from two different cohorts. The first was a group of children with type 1 diabetes (T1D) as described in [7]. Briefly, in 25 children, 6–14 years of age, with T1D across 3 sites, we acquired T1w MRI and axial 2-dimensional CSI at the level of the basal ganglia.

The second was a cohort of adults recruited in the community by word-of-mouth and flyers to study reading ability (RA). It consisted of 32 participants: 22 biologically female and 9 biologically male with a mean age of 24.0 years and standard deviation of 4.23 years. 25 were right-handed and 4 were left as measured by the Edinburgh Handedness Inventory while 3 had missing data [13]. RA was assessed with the Woodcock-Johnson IV Word Attack (WA) and Letter Word Identification (LWI) with mean scores ± standard deviation of 101.0417 ± 10.23448 and 105 ± 7.543515, respectively [14], [15]. 6 participants did not have scores available. For each participant, we then acquired SVS with and without water suppression in the left superior temporal gyrus (STG), angular gyrus (AG), and inferior frontal gyrus (IFG) with TE=30ms and a voxel size of 20x20x20mm at 3T on a Philips Achieva scanner. This produced a set of multifocal SVS data for further workflow characterization.

## 3 RESULTS

### 3.1 Spatiospectral preprocessing and quality assurance of CSI

MRS spectra are notoriously noisy, especially in CSI [6]. Additionally, the quality of spectra for assessing water-soluble metabolites can be impacted by other structures in the brain, namely dural lipids which can produce large spectral distortions under 2.0 ppm. Using this framework, we demonstrate how spatiospectral processing allowed the identification and removal of CSI voxels affected by lipid interference as described by [7] in the T1D cohort.

First, the spectra in CSI voxels with >50% coverage by the brain as segmented on T1w MRI were identified. The coverage estimates were obtained by regridding the CSI images to match the T1w MRI leveraging existing NIFTI tools and the orientation information contained in the headers. The original CSI voxels were then iterated over to identify those with majority overlap with a brain mask computed with the spatially localized network atlas tiles (SLANT) deep learning framework [16].

The resultant spectra for a representative sample were then plotted in Figure 3a. It can be seen that around 1.5 ppm two voxels contain suppressed spectra with lipid interference. To identify and remove these automatically, we fit each spectrum from the voxels across the brain with a line of best fit between 0.2 and 1.8 ppm and computed the associated root-mean-squared error (RMSE). Plotting these RMSEs, we identified the outliers using the interquartile method (Figure 5) [17]. These correspond to left frontal voxels highlighted in red. To do so, we used existing MATLAB, Python, and FSLeyes, a popular NIFTI visualization software, without special MRS-specific tools [9]. As such, we demonstrate that this framework facilitates spatiospectral preprocessing and quality assurance using existing neuroimaging standards.

**Figure 5.**
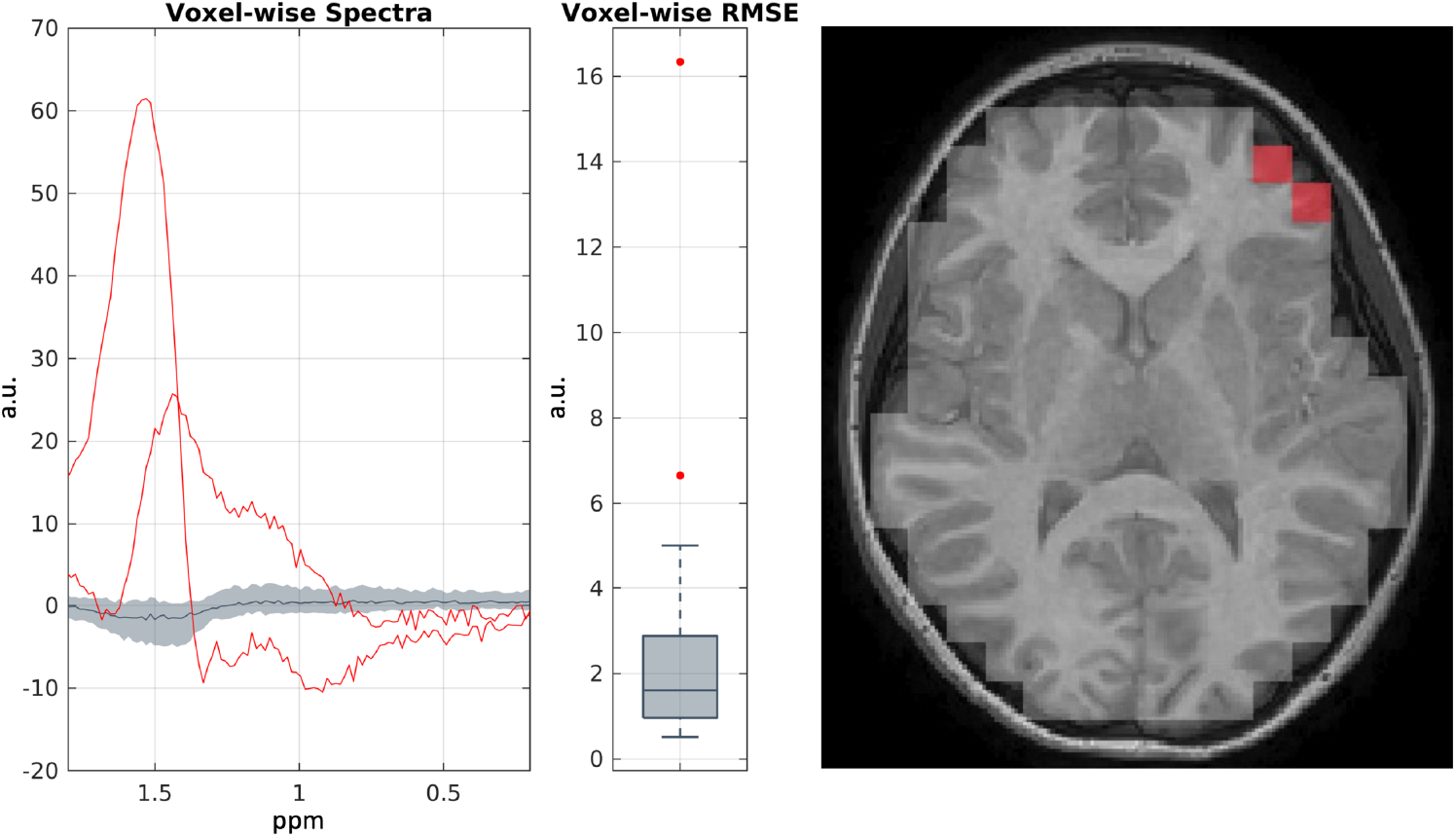
Spatiospectral preprocessing and quality assurance of CSI in a representative sample. Plotting the interquartile spectra across the brain in gray and outliers in red, we can leverage the spatial information to identify outlying spectra and exclude those voxels from further analysis. a.u. = arbitrary units.

### 3.2 Spatiospectral inference in CSI

After preprocessing and quality assurance, spatiospectral image processing can be extended to make biological inferences. We use the spatial overlap of the CSI grid with tissue segmentations performed on T1w MRI to facilitate the identification of metabolites weighted by each tissue across multiple imaging sites in the T1D cohort. The full study details are as described in [7].

Briefly, to accomplish this we use an analogous approach to the brain mask method detailed in the preceding section. We segmented the white matter (WM) and deep gray matter (dGM) nuclei using the SLANT framework (Figure 6a). Subsequently, we used the spatial information captured in the NIFTI headers for both the T1w MRI and CSI to regrid the latter to the former and computed the percent overlap of the MRS voxels to the WM and dGM segmentations. These overlap estimates were subsequently normalized across all voxels to obtain WM and dGM “weights” for each CSI voxel (Figure 6b). Using these weights, we obtained a weighted-average WM and dGM signal from the constituent CSI voxels. After this spatial processing step, spectral processing with LCModel was performed to obtain metabolite estimates in our study population in each of the tissues [5]. This approach demonstrated significant baseline differences (Figure 6c) in the ratio of the NAA peak to Cr peak (NAA/Cr), a marker of neuronal loss when reduced, between WM and dGM in our population. Thus, these results illustrate how the proposed workflow facilitates the discovery of novel spatiospectral relationships in MRS in conjunction with other structural MRI modalities.

**Figure 6.**
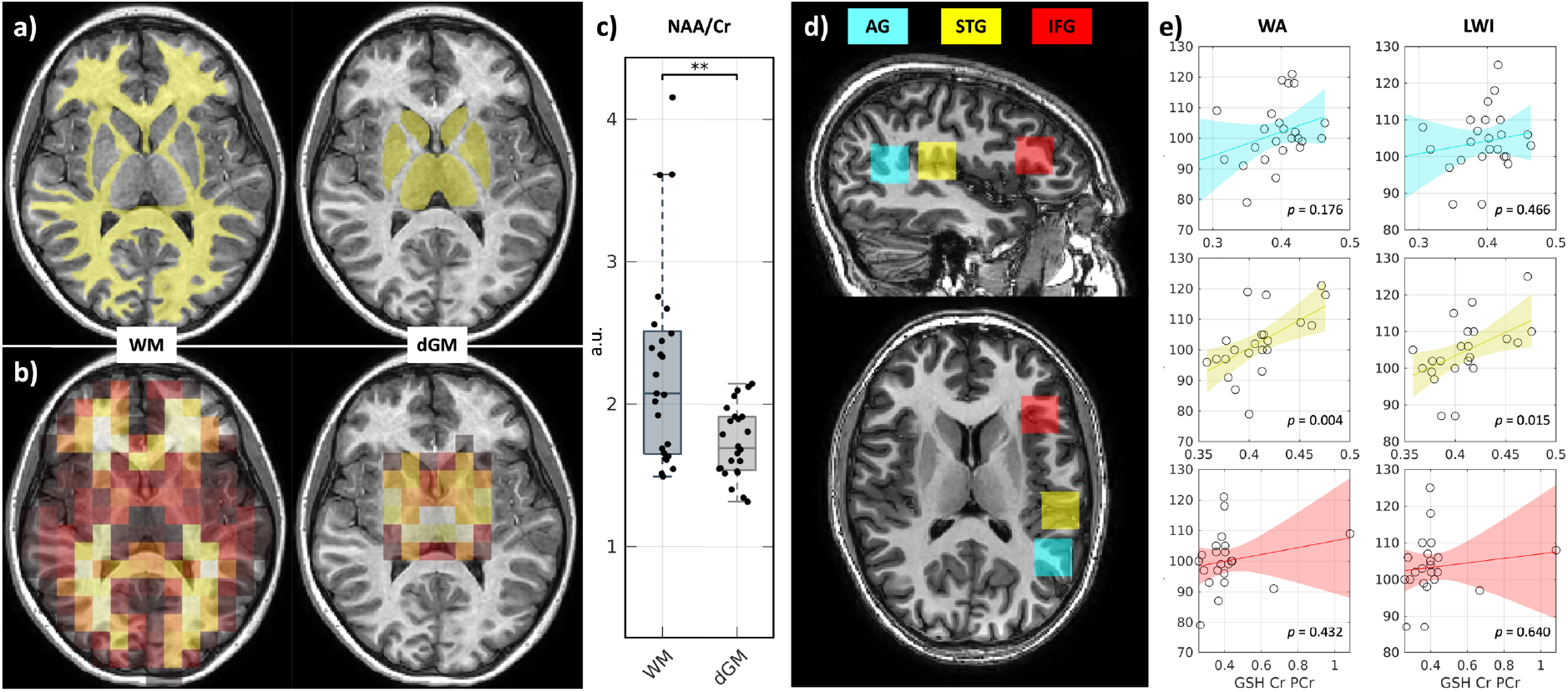
Spatiospectral inference. Using the spatial information in CSI and T1w MRI segmentations (a) captured by the proposed workflow, voxel-wise weights for consolidating spectra for each tissue are computed (b), allowing tissue-weighted spectra to be computed in this representative sample. (c) These spectra can then be spectrally processed to obtain metabolite ratios. This technique revealed baseline differences (** *p* < 0.005, Wilcoxon sign-rank test) in NAA/Cr between WM and dGM in the T1D cohort. (d) In multifocal SVS, multiple SVS locations can be measured to obtain spatial coverage. A representative sample from the RA cohort is shown. (e) Using the proposed framework, multifocal SVS studies also can be spatiospectrally processed to identify statistically significant associations of WA and LWI with neurochemical markers in different cortical gyri (*p*-values uncorrected from linear models controlling for age and handedness). a.u. = arbitrary units.

### 3.4 Spatiospectral inference in multifocal SVS

On a related note, many studies still leverage SVS over CSI. These studies utilize multiple SVS locations to obtain improved spatial coverage. We demonstrate the applicability of the proposed workflow for these studies investigating metabolic changes in the different cortical gyri captured in the RA cohort.

The time-domain SVS signals collected in each participant in each location were converted to water-suppressed real frequency spectra with FFT between 0.2 and 4.0 ppm in NIFTI format and baseline corrected with LCModel [5]. TE-specific basis sets provided by LCModel were then used to identify metabolite peaks and their associated Cramer-Rao lower bound quality estimates (%SDs). %SD was used to exclude low-quality peaks from further analysis, defined as peaks with %SD above 20 in line with LCModel recommendations. After quality control, we computed the glutathione (GSH) to Cr ratio (GSH/Cr) in each participant in each gyrus as GSH has been thought to contribute to cognition among other neuropsychiatric phenomena [18].

Using the proposed workflow in a representative RA sample, we first visualized the three SVS locations using FSLeyes (Figure 6d) [9]. To assess associations of GSH/Cr with RA in each region, we then performed linear regression of the WA and LWI scores against GSH/Cr across the cohort while controlling for age and handedness. We present the regressions with 95% confidence intervals and *p*-values for the GSH/Cr term without multiple comparisons correction in Figure 6e. In doing so, we find statistically significant associations between GSH/Cr and RA in the STG but not in the AG or IFG.

## 4 DISCUSSION

In this work, we describe a new spin on existing MRI workflows to promote spatiospectral image processing of MRS. We describe how such a workflow, building on existing neuroimaging standards, promotes flexible spatiospectral preprocessing and quality assurance of MRS as well as CSI- and SVS-based biological inference.

We note that such a workflow is not without limitations. First, as can be seen in Figure 3, different scanners use different gain for their signals, indicating that arbitration of spectral magnitude needs to be taken into consideration when performing multi-scanner studies. A similar limitation exists for MRI analysis, but the improved resolution of MRI allows for normalization of intensities to different tissues, like the cerebrospinal fluid, for instance. With low-resolution MRS, one traditional approach is to use metabolite ratios, as described herein (i.e., NAA/Cr and GSH/Cr). However, with the introduction of this workflow, new spatially-based normalization schemes can be, and should be, investigated.

Additionally, we note that a reliable spatiospectral framework opens the door for the development of new image processing routines for MRS and especially CSI. For instance, CSI suffers from known spatial biases, distortions, and nonlinearities as well as lower signal-to-noise ratios than SVS and interscanner variability [6], [7]. We hope that the introduction of this framework will spur image processing innovation in this field to overcome these challenges for more robust imaging and subsequent biological inference.

Finally, we note that making such a workflow viable relies significantly on the robust conversion of MRS DICOM to NIFTI for spatiospectral processing and NIFTI to LCModel for metabolite computation. For instance, as mentioned previously, these tools should make clear which spectra, water-suppressed or -unsuppressed, are being handled and also be verified in terms of spatial orientation with phantom studies. To facilitate the innovation of such tools in the field, we release our implementations for Philips CSI and SVS and Siemens CSI at github.com/MASILab/masimrs to augment current efforts [11].

## ACKNOWLEDGEMENTS

This work was conducted in part using the resources of the Advanced Computing Center for Research and Education at Vanderbilt University, Nashville, TN. This work was supported by the National Institutes of Health (NIH) under award numbers 1U34DK123895-01, U34DK123894-01, P50HD103537, U54HD083211, U54HD083211-S1, and T32GM007347 and by the National Science Foundation (NSF) under award number 2040462. This research was conducted with the support from the Intramural Research Program of the National Institute on Aging of the NIH. The content is solely the responsibility of the authors and does not necessarily represent the official views of the NIH or NSF.

